# Paralogues of the *PXY* and *ER* receptor kinases enforce radial patterning in plant vascular tissue

**DOI:** 10.1101/357244

**Authors:** Ning Wang, Kristine S. Bagdassarian, Rebecca E. Doherty, Xiao Y. Wang, Johannes T. Kroon, Wei Wang, Ian H. Jermyn, Simon R. Turner, J. Peter Etchells

## Abstract

Plant cell walls do not allow cells to migrate, thus plant growth and development is entirely the consequence of changes to cell division and cell elongation. Where tissues are arranged in concentric rings, expansion of inner tissue, such as that which occurs during vascular development, must be coordinated with cell division and/or expansion of the outer tissue layers, endodermis, cortex, and epidermis, in order for tissue integrity to be maintained. Little is known of how coordination between cell layers occurs, but non-cell autonomous signalling could provide an explanation. Endodermis-derived EPIDERMAL PATTERNING FACTOR-LIKE (EPFL) ligands have been shown to signal to the ERECTA (ER) receptor kinase present in the phloem. *ER* interacts with *PHLOEM INTERCALLATED WITH XYLEM* (*PXY*), a receptor present in the procambium. The PXY ligand, TRACHEARY ELEMENT DIFFERENTIATION INHIBITORY FACTOR (TDIF) is derived from *CLE41* which is expressed in the phloem. These factors therefore represent a mechanism by which intertissue signalling could occur to control radial expansion between vascular and non-vascular tissue in plant stems. Here we show that *ER* regulates expression of *PXY* paralogues, *PXL1* and *PXL2*, and that in turn *PXY*, *PXL1* and *PXL2* together with *ER*, regulate the expression of *ERL1* and *ERL2*, genes paralogous to *ER*. *PXY*, *PXL1*, *PXL2* and *ER* also regulate the expression of ER-ligands. Genetic analysis of these six receptor kinase genes demonstrated that they are required to control organisation, proliferation and cell size across multiple tissue layers. Taken together, our experiments demonstrate that ER signalling attenuates *PXL* expression in the stem, thus influencing vascular expansion and patterning. We anticipate that similar regulatory relationships, where tissue growth is controlled via cell signals moving across different tissue layers, will coordinate tissue layer expansion throughout the plant body.

## Introduction

Cell migration is fundamental to development of animal body plans. By contrast, plant cell walls do not allow cells to migrate and consequently plant growth and development is entirely a result of differential growth. As such, initiation and elaboration of plant organs occurs via coordinated changes to the orientation and occurrence of cell divisions, and by cell expansion. In embryos, pattern is established early in development. 28-cell embryos have already specified the provascular tissue which consists of four cells the centre of the embryo, a layer of endodermal tissue which surrounds the provasculature, and an outer layer of epidermal cells (ten Hove et al., 2015). Extra tissue types are subsequently specified via specific rounds of asymmetric cell divisions (Kajala et al., 2014) such that in the dicot hypocotyl, the tissue pattern along the radial axis is epidermis-cortex-endodermis-pericycle-phloem-cambium-xylem. The hypocotyl maintains a similar pattern throughout the life of the plant (Chaffey et al., 2002), with the exception of the epidermis and cortex, which are replaced by periderm as the hypocotyl expands (Wunderling et al., 2018). Thus, coordination of tissue expansion must occur as organs increase in size. This occurs both at the level of cell division, where cell number increases from tens to hundreds to thousands of cells, and at the level of cell size, which in differentiated cells differs according to function.

Little is known about how patterns are maintained through very large increases in plant size. However, evidence points to the presence of mechanisms that coordinate the order of tissue layers. In the Arabidopsis root, removal of the root tip results in a reorganisation of the organ to enable the formation of a new meristem. Strikingly, stable patterning of tissue layers is established in the reorganised tissue separately from the activity of the stem cell niche. This suggests that tissue layer organisation is independent of stem cell growth (Efroni et al., 2016). Non-cell autonomous signalling represents one mechanism through which tissue layer organisation could be coordinated. A ligand secreted by one tissue could provide positional information to a receptor located in an adjacent cell type. Ligand-receptor pairs that signal between tissue layers and are required for tissue layer organisation have been described. For example, in microsporangial patterning, TAPETUM DETERMINANT1 (TPD1) ligand is excreted from microsporocytes and perceived by the EXCESS MICROSPOROCYTES 1 (EMS1) receptor, present in adjacent locular peripheral cells which, in turn, go on to form the tapetum (Jia et al., 2008). The consequences of disrupting this interaction includes disruption of the integrity of the tapetal cell layer and changes to the orientation of cell division (Feng and Dickinson, 2010).

In vascular development, spatially separate ligands and receptors are also required to regulate vascular organisation. TRACHEARY ELEMENT DIFFERENTIATION INHIBITORY FACTOR (TDIF) ligand is encoded for by three genes, *CLAVATA3-LIKE/ESR-RELATED 41* (*CLE41*), *CLE42* and *CLE44.* It is excreted from the phloem and perceived by the PHLOEM INTERCALLATED WITH XYLEM/TDIF receptor (PXY/TDR) receptor, which is cambium-expressed. The consequence of loss of the TDIF-PXY interaction is loss of organised tissue layers characterised by disruption to the spatial separation of xylem, cambium, and phloem. Loss of PXY also results in reductions in cell division in the cambium, and premature xylem differentiation (Etchells and Turner, 2010; Fisher and Turner, 2007; Hirakawa et al., 2008; Ito et al., 2006). TDIF-PXY interacts with a second ligand-receptor pair to maintain the spatial separation of tissues in the vasculature. In stems, the *ERECTA* (*ER*) receptor is expressed in the phloem, and its cognate ligands, *CHALLAH-LIKE 2*/*EPIDERMAL PATTERNING FACTOR-LIKE 4* (*CLL2*/*EPFL4*) and *CHALLAH* (*CHAL/EPFL6*) are expressed in the endodermis (Abrash et al., 2011; Uchida et al., 2012). *pxy er* mutant stems show organisation defects greater than those of *pxy* single mutants (Etchells et al., 2013). Thus the genetic interaction between EPFL-ER and TDIF-PXY represents a non-cell autonomous signalling system that organises tissue layers between endodermis, phloem, cambium and xylem (Figure 1B). In hypocotyls, *ER* expression is reported to be much broader (Ikematsu et al., 2017), but nevertheless changes to organisation of vascular tissues in *er pxy* hypocotyls are also apparent (Etchells et al., 2013).

**Figure 1.**
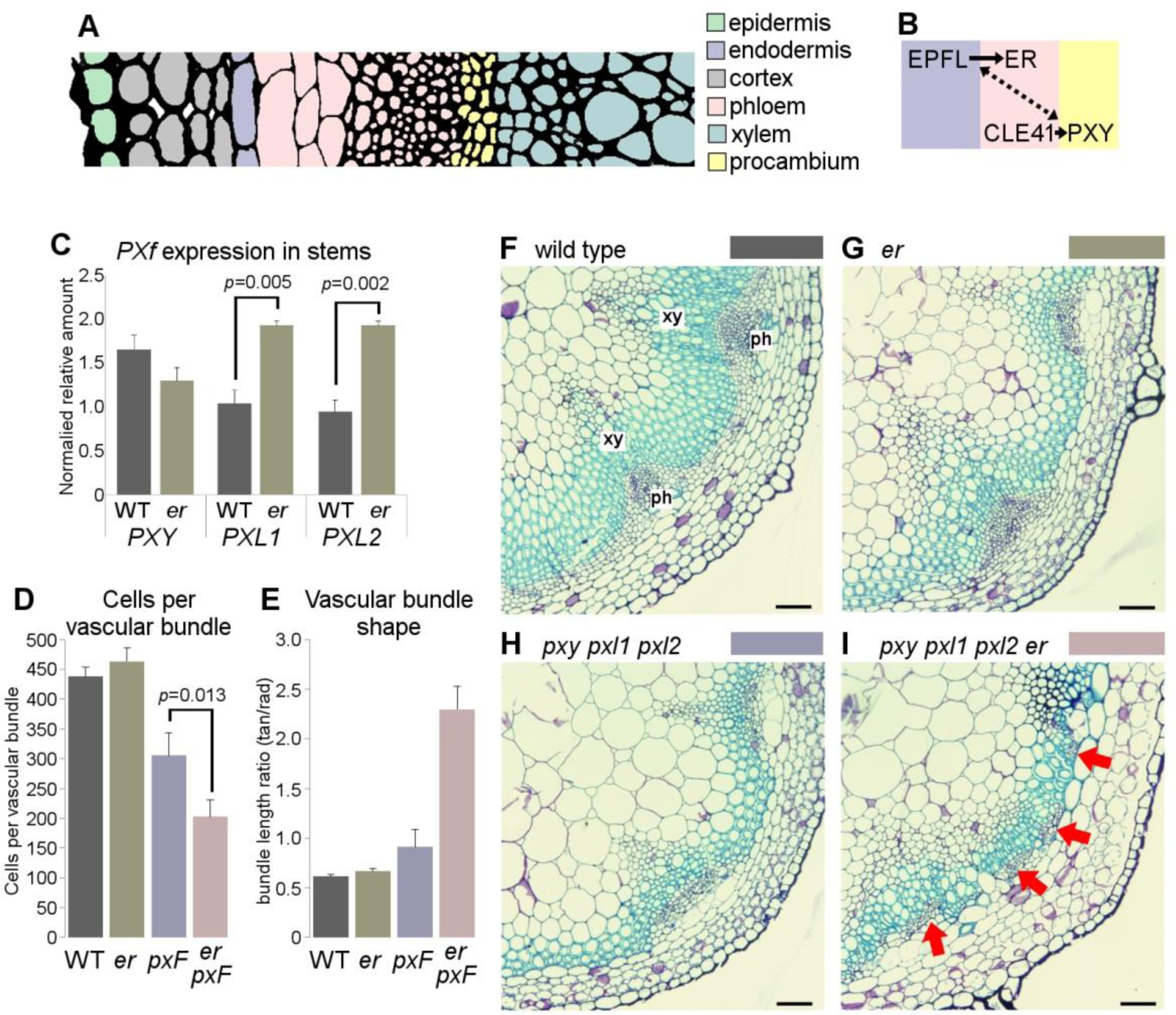
Interaction between *PX*f and *ER*. (A) Tissue types in the Arabidopsis stem. (B) Interactions between PXY and ER signalling are characterised by non-cell autonomous interactions. (C) qRT-PCR showing elevated expression of *PXL1* and *PXL2* in *er* mutants (expression normalised to *ACT2*). (D) Graph showing mean cells per vascular bundle. (E) Representation of vascular bundle arrangement (ratio of size along tangential/radial axes). (F-I) Transverse sections through wild type (F), *er* (G), *px*f (H) and *px*f *er* (I) stems. Arrows in (I) point to phloem distributed around the stem, rather than in discrete bundles. *p* values were calculated using a student’s t-test (C), or ANOVA with an LSD post-hoc test (D). Scales (F-I) are 50 µM. xy marks xylem; ph marks phloem.

In the Arabidopsis genome, both *PXY* and *ER* paralogues are present. The *PXY* family, hereafter referred to as *PX*f is constituted of *PXY*, *PXY-LIKE1* (*PXL1*), and *PXY-LIKE2* (*PXL2*). *pxl1* and *pxl2* enhance the vascular organisation defects that are characteristic of *pxy* mutants (Etchells et al., 2013; Fisher and Turner, 2007). The *ER* paralogues are *ER-LIKE1* (*ERL1*) and *ERL2* (Shpak et al., 2004). The *ERECTA* family (*ER*f) have wide ranging roles in regulation of plant growth and development. Redundantly, these three genes function in cell elongation, cell division, inflorescence architecture (Shpak et al., 2004; Torii et al., 1996), floral patterning (Bemis et al., 2013), shoot apical meristem fate (Kimura et al., 2018; Uchida et al., 2013), and stomatal spacing (Shpak et al., 2005). In the context of plant vascular development, they regulate vascular expansion in the stem (Uchida and Tasaka, 2013) and hypocotyl where they control the timing of xylem fibre formation, and levels of radial growth (Ikematsu et al., 2017; Ragni et al., 2011). A hallmark of loss of *ER*f genes is an increase in cell size, particularly with respect to the radial axis (Shpak et al., 2004; Shpak et al., 2003).

In this paper we investigated the coordination between tissue layers required for plant growth and development. In particular we examined the mechanism by which *PX*Y and *ER* signalling coordinates development between vascular and non-vascular cell types. We found that in *Arabidopsis*, *ER* together with members of the *PX*f family coordinate the expression of *ERL1, ERL2*, *EPFL4* and *EPFL6*. The observation that vascular-expressed *PXY* and *ER* regulate the expression of non-vascular-expressed *EPFL4* and *EPFL6* in stems, demonstrates coordination of growth regulators between vascular and non-vascular tissue layers. To understand *ER*f function in the context of *PXY* signalling, we generated *pxy pxl1 pxl2 er erl1 erl2* sextuple mutants using a combination of classical genetics and genome editing. Analysis of these lines demonstrated that the *PX*f and *ER*f interact to coordinate tissue integrity at the levels of both cell size, and cell division. Our results demonstrate that *PX*f and *ER* together control tissue growth via coordination of gene expression between vascular and non-vascular cell types.

## Materials and Methods

### Accession numbers

AGI accession numbers for the genes studies in this manuscript: At3g24770 (*CLE41*), At5g61480 (*PXY*), At1g08590 (*PXL1*), At4g28650 (*PXL2*), At2g26330 (*ER*), At5g62230 (*ERL1*), At5g07180 (*ERL2*), At4g14723 (*CLL2*/*EPFL4*), At3g22820 (*CLL1*/*EPFL5*), At2g30370 (*CHAL*/*EPFL6*).

### Gene expression

For qRT-PCR, RNA was isolated using Trizol reagent (life technologies) prior to DNAse treatment with RQ1 (promega). cDNA synthesis was performed using Tetro reverse transcriptase (Bioloine). All samples were measured in technical triplicates on biological triplicates. qPCR reactions were performed using qPCRBIO SyGreen Mix (PCR Biosystems) using a CFX connect real time system (Bio-Rad) with the standard sybr green detection programme. A melting curve was produced at the end of every experiment to ensure that only single products were formed. Gene expression was determined using a version of the comparative threshold cycle (Ct) method using average amplification efficiencies of each target as determined using LinReg PCR software (Ramakers et al.,2003)). Samples were normalised to *18S* rRNA or *ACT2*. Primers for qRT-PCR are described in Table S1. Significant differences in gene expression were identified with ANOVA and LSD post-hoc test.

### Plant lines

Previously described parental lines *pxy*-*3 pxl1-1 pxl2-1* (referred to hereafter as *er*f) and *pxy-5 er-124* (Etchells et al., 2013) were crossed to generate *pxy*-*3 pxl1-1 pxl2-1 er-124* (*er px*f). The quadruple mutants were selected in the F3 by PCR using primers listed in Table S1. To generate *px*F *er erl2* quintuple mutants, parental lines *erl1, er-105 erl1-2/+ erl2-1* (Shpak et al., 2004) and *pxy*-*3 pxl1-1 pxl2-1* (Etchells et al., 2013) were crossed. Plants homozygous for *er* were selected by visual phenotype in the F2, which was also sprayed with glufosinate to select for plants carrying an *erl2-1* allele. Families homozygous for glufosinate resistance in the F3 were screened for *pxy-3*, *pxl1-1* and *pxl2-1* to generate *px*f *er erl2*. *er* and *erl2* mutants were subsequently confirmed by PCR.

*erl1* genome edited lines were generated using an egg cell specific CRISPR/Cas9 construct (Wang et al., 2015; Xing et al., 2014). Briefly, target sequences TCCAATTGCAGAGACTTGCAAGG and TCTTGCTGGCAATCATCTAACGG were identified using the CRISPR-PLANT website (Xie et al., 2014) and tested for off-targets (Bae et al., 2014). Primers incorporating the target sequences (Table S1) were used in a PCR reaction with plasmid pCBC-DT1T2 as template to generate a PCR product incorporating a guide RNA against *ERL1*. A golden gate reaction was use to incorporate the purified PCR product into pHEE2E-TRI. The resultant *ERL1* CRISPR/cas clone was transferred to Arabidopsis by floral dip (Clough and Bent, 1998). *erl1*^GE^ mutants were selected in the T1 generation by sequencing PCR products generated from primers specific to ERL1 genomic DNA that flanked the guide RNA target sites.

For spatial expression of *ER*f genes in *pxy* or *er*, previously described *ER::GUS*, *ERL1::GUS* and *ERL2::GUS* reporters were used (Shpak et al., 2004) which were crossed to *pxy-3* or *er-124*. *pxy* mutants were selected in the F2 using primers described Table S1. Reporter lines were picked which also demonstrated GUS expression as judged by *GUS* histochemical staining, and the presence of *GUS* reporter construct was subsequently confirmed by PCR using primers described in Table S1.

*Nicotiana benthamina* lines overexpressing *AtCLE41* were generated by transforming a previously described *35S::AtCLE41* binary plasmid (Etchells and Turner, 2010) using the method described by Horsh (Horsch et al., 1985).

### Analysis of vascular tissue anatomy

Vascular morphology was assessed using tissue embedded in JB4 resin. For vascular bundles, inflorescence stem tissue from 0.5 cm above the rosette was assessed. Tissue was fixed in FAA, dehydrated in ethanol and infiltrated with JB4 infiltration medium, prior to embedding. 4 µM sections, taken using a Thermo Fisher Scientific Finesse ME 240 microtome were stained in 0.02% aqueous toluidine blue and mounted with histomount.

GUS stained tissue was harvested to cold phosphate buffer on ice. Samples were treated with ice-cold acetone for 5 minutes and then returned to phosphate buffer. GUS staining buffer (50 mM phosphate buffer, 0.2% triton, 2 mM potassium ferrocyanide, 2 mM potassium ferricyanide, 2 mM X-Gluc) was added and samples were infiltrated using a vacuum, before incubation overnight at 37°C. Samples were progressively incubated in: FAA, 70%, 85%, 95% EtOH for 30 minutes each prior to embedding in Technovit 7100 according to the manufacturer’s instructions. Embedded samples were allowed to polymerize at room temperature for two hours and at 37◦C overnight until solid.

The inhibition layer was removed by wiping with a lint-free cloth. Samples were sectioned, counter-stained with 0.1% neutral red and mounted using histomount.

### Quantitative morphology calculations

Images of five-week old wild type *px*F and *px*F *er erl2* hypocotyls were used. Images from 6 different individuals were selected for each genotype tested. From each image, a minimum of 10 cells of each cell type (xylem vessels, xylem fibers, phloem and parenchyma) were selected from a wedge with a 60 degree central angle (Figure S1A). A MATLAB code was generated to extract the intrinsic properties of each cell type. To that end, the code was designed to split each image into binary sub-images, wherein the interior of the cell type of interest was represented as white objects on black background (Figure S1B). The cells from each image were then analysed as connected components of the image and their area, perimeter and ellipticity (calculated as the ratio of major to minor axis) extracted. To remove noise, i.e. data obtained from objects which were wrongly classified as connected components within the algorithm (e.g. stray pixels), the code was devised to discard data outside the range of pre-specified number of standard deviations from the mean. The number of standard deviations differed between the different genotypes and was selected on a trial-and-error basis, by referring back to the original images, with the objective of maintaining the maximum number of viable data points (2-3 standard deviations for xylem cells, 1.4-1.6 standard deviations for fibre cells, 3 standard deviations for all other cell types). The data was converted from pixels to microns using a calibration factor, in order to yield results consistent with laboratory observations.

To test the significance of the variation between the cell areas and perimeters between the different genotypes, a Lilliefors test was performed which determined that the data was normally distributed at the 5% significance level, allowing for the subsequent use of a nested ANOVA in R. To perform the nested ANOVA, the data was classified according to the treatment (i.e. genotype) and plant ID within that treatment, with the response variable either the area or perimeter. Following the results of the nested ANOVA, a post-hoc Tukey HSD test was performed to determine the significance of the pairwise differences between the means of the areas/perimeters within each genotype. Due to the varying number of cells for each genotype, the histogram and boxplot data representations were derived from a random sample for each cell type, where a maximum number of representatives from each genotype were selected. For each genotype, the MATLAB code was designed to randomly select 70 xylem cells, 340 fibre cells, 200 phloem cells, 320 parenchyma cells.

Mean hypocotyl area was calculated from images of six plants of each genotype. A MATLAB code was used to measure the length of the shorter and longer radius from each image (one radius at 12 or 6 o’clock and one radius at 3 or 9 o’clock, as appropriate). The length of the radii in pixels was subsequently converted to microns and the formula for the area of ellipse (*A* = *r*_1_ ∗ *r*_2_ ∗ π) used to calculate the area of each hypocotyl. A Lilliefors test at 5% significance level was used to confirm that the areas for each genotype were normally distributed. A one-way ANOVA was performed to establish the existence of significant variation between the areas of the different genotypes, followed by a post-hoc Tukey HSD test to gain insight into the pairwise variation between the means.

## Results

### PXL1 and PXL2 are regulated by ER in the stem but not in the hypocotyl

*pxy* mutants demonstrate radial patterning defects including intercalation of vascular cell types (Fisher and Turner, 2007). These defects are enhanced by mutations at the *ER* locus (Etchells et al.,2013). ER ligands, EPFL4 and EPFL6 (Abrash et al., 2011), are expressed outside of the vascular cylinder, in the endodermis. *ER* is expressed in the phloem (Uchida et al., 2012). TDIF encoding genes, *CLE41*, *CLE42* and *CLE44*, are expressed in the phloem, and TDIF signals to PXY which is expressed in the procambium (Etchells and Turner, 2010; Fisher and Turner, 2007; Hirakawa et al., 2008). Therefore, *PXY* and *ER* constitute a mechanism by which coordination of radial expansion between vascular and non-vascular tissues could occur (Figure 1A-B). To explore the interaction between *PXY* and *ER*, we sought to determine whether ER signalling could regulate PXY signalling and/or vice-versa at the level of gene expression. We have previously shown that *ER* does not regulate expression of *CLE41*, *CLE42* and *CLE44* (Etchells et al., 2013). Consequently we asked whether *ER* might regulate expression of the *PX*f family of receptors. qRT-PCR was used to test levels of *PX*f gene expression in stems and hypocotyls of wild type and *er* in 5 week old plants. In hypocotyls, the level of *PX*f gene expression was unchanged in *er* mutants compared to wild type (Figure S2A). By contrast, *PXL1* and *PXL2* expression, but not that of *PXY* was found to be elevated in *er* mutant stems (Figure 1C). These observations suggest that *ER* signalling may regulate vascular development by setting *PXL1* and *PXL2* levels in the stem. To determine the function of *PXL1* and *PXL2* regulation by *ER*, specifically in the context of the *ER-PXY* interaction, *er px*F quadruple mutants (*er pxy pxl1 pxl2*) were generated.

In previous studies, when *px*f plants were compared to *pxy* single mutants, vascular organisation defects were enhanced (Fisher and Turner, 2007), but hypocotyl radial expansion and the number of cells per vascular bundle did not differ between *pxy* and *px*f (Etchells et al., 2013). Here, in inflorescence stems, *er px*f lines had considerably fewer cells per vascular bundle than either *px*f or *er* counterparts (Figure 1D; Table S2). Therefore *PXL1* and *PXL2* do function redundantly with *ER* to regulate vascular proliferation. Furthermore, a similar reduction in proliferation was observed in the hypocotyl. Here, although *PXL1* and *PXL2* expression was unaffected in *er* mutants (Figure S2), the diameter of *er px*f quadruple mutant hypocotyls was nevertheless significantly smaller than controls (Figure S2B; Table S2).

While changes to vascular proliferation were apparent in *er px*f inflorescence stems, by far the most dramatic defect was observed in when vascular bundle shape was assessed (Figure 1D-E). Typically in *Arabidopsis* stems the distribution of vascular bundles is such that there is a greater distribution of vascular tissue along radial axis of the stem than along the tangential. We measured the ratio of tangential:radial length of wild type vascular bundles to be 0.61, and in *px*F lines this ratio rose to 0.91 (Figure 1F-I; Table S2). We have previously shown that in *pxy er* stems this ratio is 1.36 (Etchells et al., 2013). In *er px*f stems a dramatic redistribution of vascular cell types along the radial axis was observed when compared to all other lines tested, such that the ratio of tangential:radial length of vascular tissue was 2.30 (Figure 1E-I; Table S2). In some plant stems this led to an almost complete ring of vascular tissue, with phloem cells scattered around the circumference of the vascular cylinder (arrows in figure 1I), rather than present in discrete vascular bundles. Thus *PXL1* and *PXL2* are critical in regulating radial pattern, particularly in the absence of *ER* and *PXY*, and these data support the idea that *ER* and *PX*f constitute a mechanism for organising layers of cell types within the vasculature.

### Co-regulation of *ER*f expression by ER and PXL

Having observed that *PX*f genes were differentially expressed in *er* mutants, and that *PXL1* and *PXL2* contribute to the control of radial pattern, we also sought to determine whether members of the *ER* gene family might also be regulated by *ER*, or indeed by the *PX*f. *ER* expression levels in stems and hypocotyls of *px*f lines did not differ from wild type, as determined by qRT-PCR. Expression levels of *ERL1* and *ERL2* did not differ significantly in neither *er* mutants nor in *px*f triple mutant stems compared to wild type (Figure 2A-C). By contrast, *ERL1* expression was reduced in *er px*f lines compared to *er* single mutants. Thus *ERL1* expression is maintained by an interaction between *PX*f and *ER* in stems (Figure 2A), and as such this interaction is required to maintain the requisite level of *ER*f signalling.

**Figure 2.**
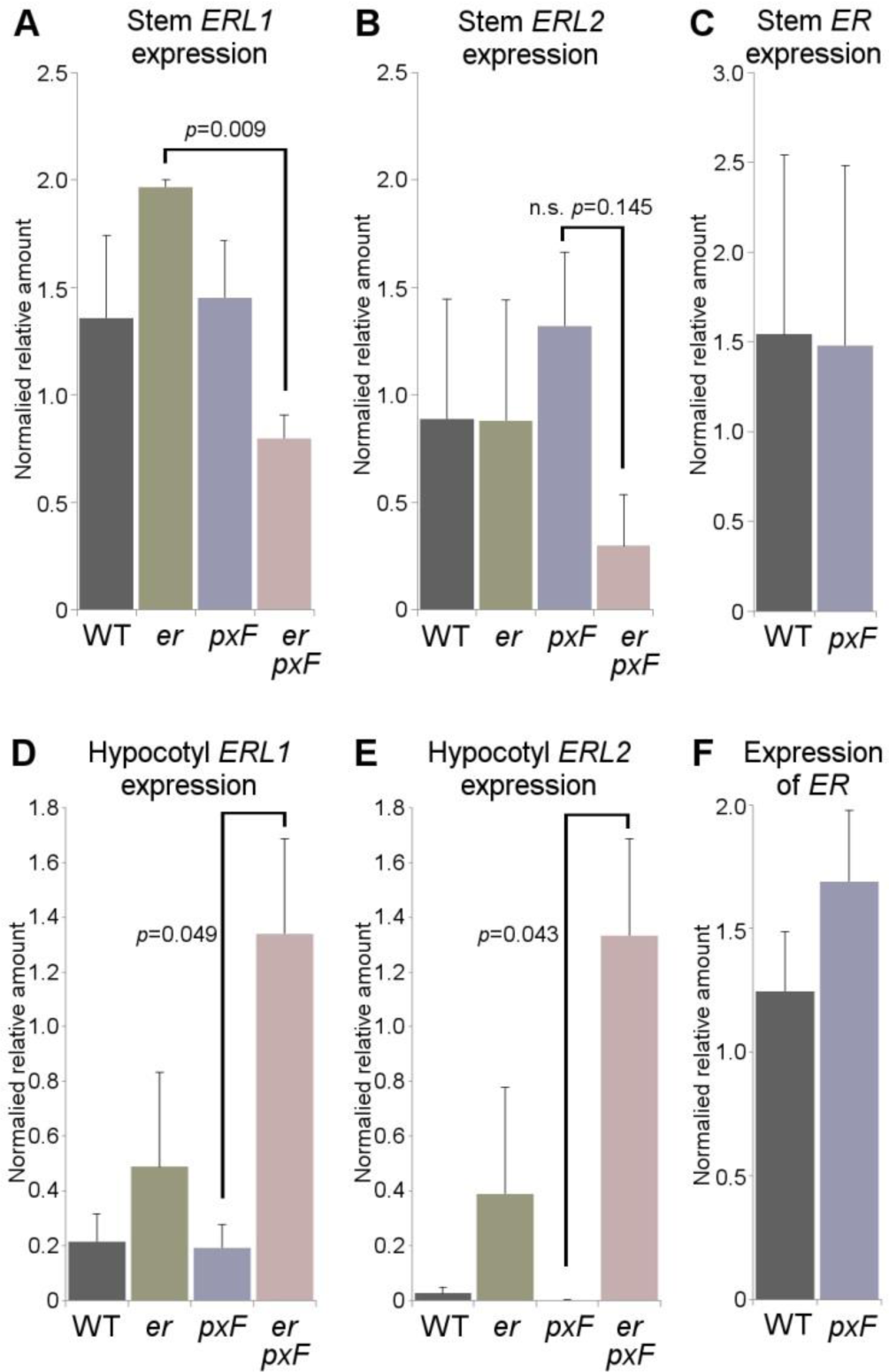
qRT-PCRs measuring *ER*f expression. (A-C) Stem expression of *ERL1* (A), *ERL2* (B) and *ER* (C) in wild type, *er*, and *px*f mutants in stems. Expression was normalised to 18S rRNA. (D-F) Expression of *ERL1* (D), *ERL2* (E) and *ER* (F) in hypocotyls (normalised to 18S rRNA). *p* values were calculated with ANOVA and LSD post-hoc test.

In hypocotyls, *ERL1* acts redundantly with *ER*, negatively regulating hypocotyl growth and the timing of xylem fibre differentiation (Ikematsu et al., 2017). *ERL2* has not been assigned a function in hypocotyl development as its expression has been reported as absent from hypocotyls in 9 day old seedlings and 3 week old plants (Ikematsu et al., 2017; Uchida et al., 2013). To understand how *PXY* and *ER* might influence *ER*f expression, *ER*f:*GUS* reporter constructs (Shpak et al., 2004) were crossed into *pxy* and *er* mutants. To our surprise, in 5 week plants we did detect *ERL2::GUS* reporter expression in the hypocotyls of wild type which, at this growth stage, demonstrated a very similar pattern to that observed for *ERL1* and *ER*. Thus, *ERL2* expression is a feature of late hypocotyl development (Figure S3). *ER*, *ERL1* and *ERL2* expression was present in most hypocotyl cell types, with two maxima; the first in the cambium and xylem initials, and the second in the cortex (Figure S3A, D, G; arrowheads). No change in the pattern of *ERL1* or *ERL2::GUS* expression was observed in *er* mutants (Figure S3C, F). However, the clearly defined expression maxima that were observed in *ER::GUS*, *ERL1::GUS* and *ERL2::GUS* lines in both wild type and *er* mutants, lacked definition in the absence of *PXY*. Here, for all three reporters expression was observed to be more even across the hypocotyl (Figure S3B, E, H), possibly due to the changes in vascular organisation in *pxy* mutants.

Having defined the pattern of *ER*f expression, at least in a subset of genotypes, we then sought to address changes to *ER*f expression levels. In common with our observation in the stem, hypocotyl *ERL1* and *ERL2* expression did not differ between wild type, *er*, and *px*f lines as determined by qRT-PCR. Our expectation was that *ER*f levels would be reduced in *px*f *er* hypocotyls, as they were in the stem (Figure 2A-C), but by contrast, a striking increase in *ERL1* and *ERL2* gene expression was observed in *px*f *er* hypocotyls (Figure 2D-F), and as such, opposite regulation of *ERL1* and *ERL2* by *ER* and *PX*f genes occurs in the hypocotyls and stem.

*EPFL4* and *EPFL6* encode the ligands that signal to ER during vascular development (Uchida and Tasaka, 2013) and stem elongation (Abrash et al., 2011; Uchida et al., 2012). *EPFL5* genetically interacts with *EPFL4* and *EPFL6* (Abrash et al., 2011), so these three genes were included in our qRT-PCR analysis. In hypocotyls, no changes were observed in *EPFL4/5/6* expression levels in *er*, *px*f or *er px*f genotypes (Figure S4D-F). However, inflorescence stem expression of *EPFL4* and *EPFL6*, but not that of *EPFL5*, demonstrated significant reductions in expression in *er px*f lines (Figure S4A-C). Thus *PX*f and *ER* interact to control *EPFL* ligand expression in addition to that of their cognate receptors, *ERL1* and *ERL2*.

### Coordination of hypocotyl size is lost in *PX*f *ER*f mutants

The *PX*f promotes radial growth in hypocotyls (Etchells et al., 2013; Fisher and Turner, 2007; Hirakawa et al., 2008), whereas *ER* and *ERL1* signalling represses it (Ikematsu et al., 2017). Thus our gene expression data demonstrating that *PX*f plays a part in repression of *ERL* expression in hypocotyls (Figure 2D-E) is consistent with existing phenotypic data where *PX*f might be expected to repress expression of negative regulators of hypocotyl radial growth. In addition to repressing radial growth, *ER* and *ERL1* have also been described as preventing premature fibre formation as *er erl1* hypocotyls develop fibre cells where parenchyma are present in wild type. *ERL2* is thought not to function in the hypocotyl given its very low expression levels in the early stages of development (Ikematsu et al., 2017). Prior to addressing the function of elevated *ERL* expression in *px*f *er* hypocotyls, we first sought to verify whether this was indeed the case as we found *ERL2* to be expressed in hypocotyls at 5 weeks (Figure S3D). We tested if *ERL2* functioned similarly to *ERL1* by analysing *er erl2* lines. Neither change to fibre formation, nor to hypocotyl radial growth were observed (Figure S5), thus in contrast to *ERL1* (Ikematsu et al., 2017), a function for *ERL2* in hypocotyl development is not apparent in an *er erl* double mutant background. To address the function of the elevated *ERL* expression that was observed in *px*f *er*, we generated and analysed *px*f *er erl2* quintuple mutants. *px*f *er erl2* hypocotyl diameters were dramatically smaller than those of parental lines (Figure 3), and therefore elevated *ERL2* expression in *px*f *er* hypocotyls is required to maintain hypocotyl growth rates.

**Figure 3.**
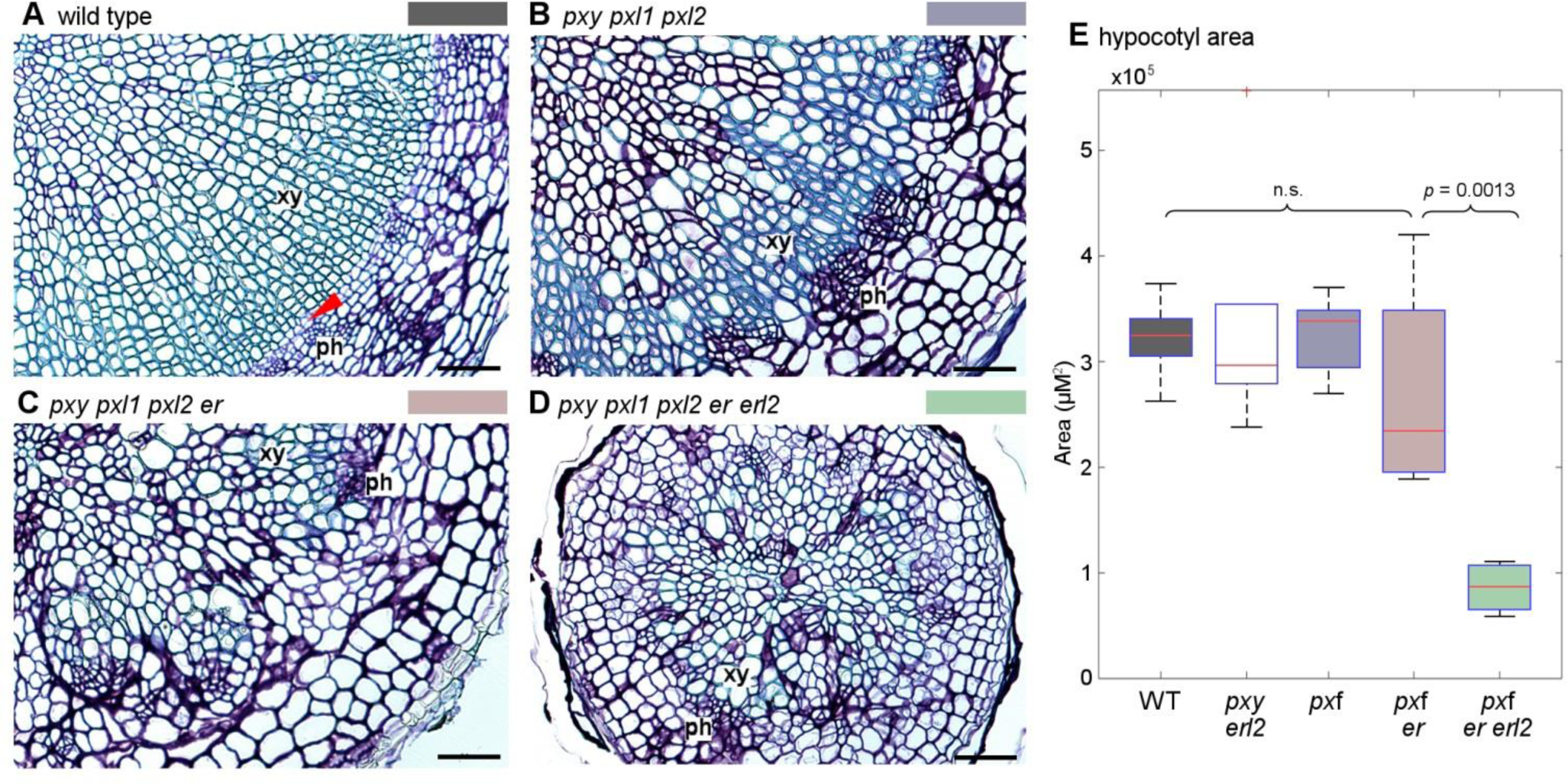
*erl2* enhances *px*f *er*. (A-D) Transverse sections through *Arabidopsis* hypocotyls. (A) Wild type. (B) *px*f. (C) *px*f *er*. (D) *px*f *er erl2*. (E) Box plot showing comparison of hypocotyl area of *px*f and *er erl2* combinatorial mutants. *p* value was calculated with ANOVA and a Tukey post-hoc test. xy is xylem; ph is phloem; red arrowhead in (A) marks dividing cambium. Scales (A-D) are 50 µM.

One common characteristic of mutants with reduced cell division is an increase in cell size, relative to wild type plants. This compensates for fewer cells, such that final organ size is often similar to that of wild-type plants (Horiguchi and Tsukaya, 2011). In the course of our analysis, cell sizes and shapes appeared to differ among our mutant lines, and in particular, cells of *px*f appeared larger than those other lines (Figure 4A, B). Consequently, cell morphology was calculated from anatomical sections (Figure S1A-B). Cell area, perimeter and ellipticity were determined for xylem vessels, fibres, parenchyma, and phloem cells in wild type, *px*f, and *px*f *er erl2* lines. Xylem vessels and fibres in *px*f lines demonstrated increases in cell area, relative to both wild type and *px*f *er erl2* plants. The average area of phloem and parenchyma cells proved not to differ significantly between all the genotypes tested (Figure 4). Xylem cells are characterised by rigid secondary cell walls, so we hypothesised that parenchyma may be subject to changes in cell shape to accommodate the increased xylem cell size. To test this hypothesis, we calculated the ellipticity, of the parenchyma and other hypocotyl cell types by determining major to minor axis ratios, but this parameter varied little between genotypes (Figure S1C-F). Finally, we measured the perimeter of each of the four cell types. In this analysis we determined that the perimeters of xylem vessels, parenchyma, and phloem cells were significantly larger in *px*f lines than in wild type (Figure 4; RHS). A near-significant difference (*p*= 0.053) was observed in the case of fibres. No differences were observed between wild type and *px*f *er erl2* lines. Therefore, in hypocotyls, *px*F mutants compensate by increasing cell size, but these cellular changes are entirely dependent on *ER* and *ERL2*. Taken together, PXY and ER signalling interact to coordinate organ size, at the levels of cell size, proliferation, and pattern maintenance.

**Figure 4.**
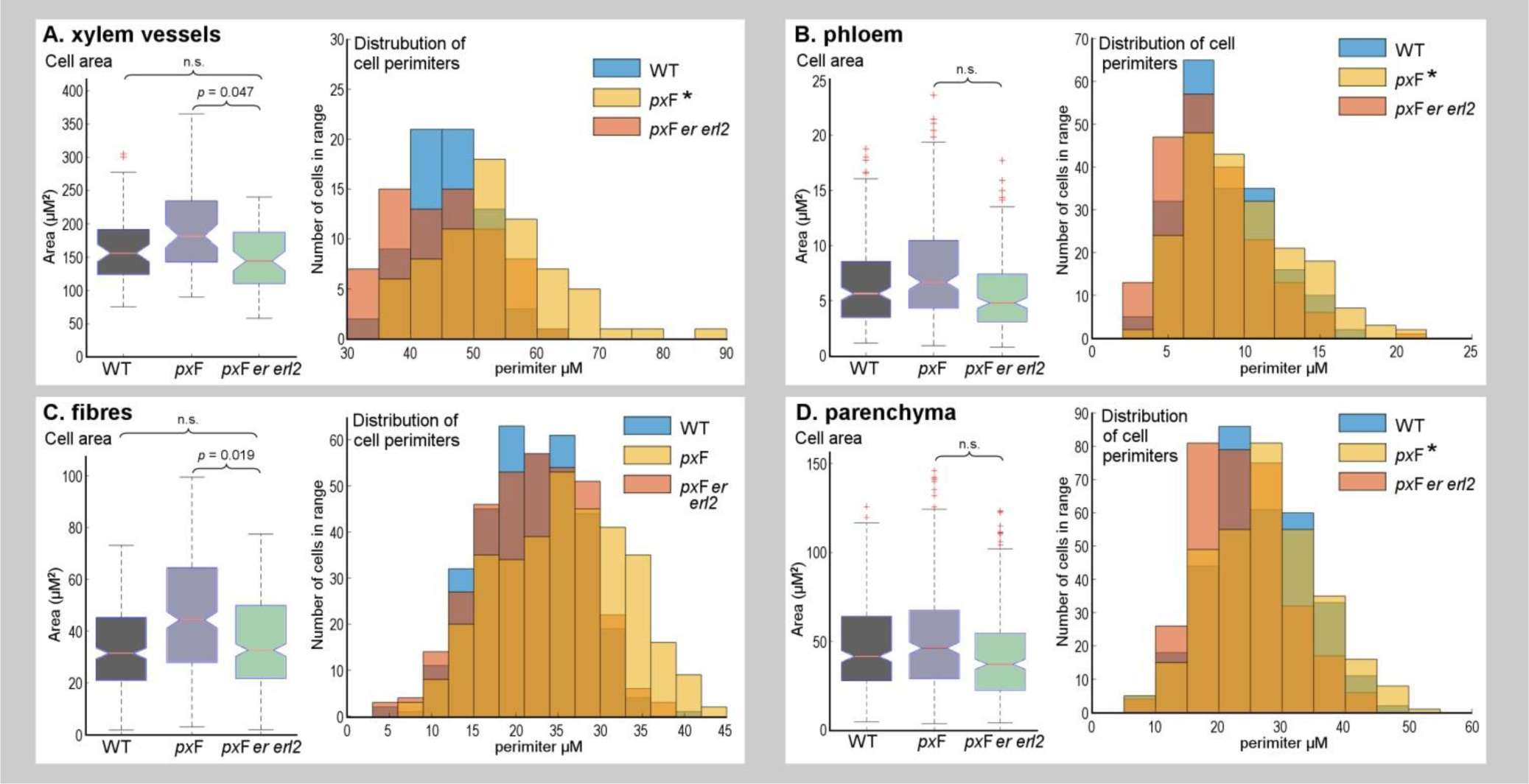
Comparisons of hypocotyl cell morphology. (A-D) Notched boxplots on left show mean area per xylem vessel (A), phloem cell (B), xylem fibres cell (C) and parenchyma cell (D). Notches show the 95% confidence level of the median. Histograms on right show the distributions of cell perimeters. Asterisks on the colour key mark where *px*f perimeters were greater than those of *px*f *er erl2* lines, i.e. xylem vessels, phloem, and parenchyma (*p* < 0.05). For fibres, *p* = 0.053. Differences were calculated with ANOVA and a Tukey post-hoc test.

### Defects of *px*f *er*f sextuple mutants

We sought to remove all *PX*f and *ER*f genes to understand any remaining redundancy between these two gene families. However, *PXY* and *ERL1* are tightly linked on chromosome 5, separated by just 270 Kb. To overcome this linkage we designed a CRISPR/cas9 construct that contained two guide RNAs against *ERL1* (Figure S6) in order to remove the remaining function *ER*f gene from *px*F *er erl2*. The resulting *px*f *er*f sextuple mutants were compared to *px*f *er erl2* lines, *er*f, *px*f, and wild type. Although gross morphology of *px*f *er*f sextuple lines was considerably smaller than *px*f *er erl2* counterparts (Figure S7), inflorescence stem vascular morphology was similar in these two lines (Figure 5). Both were characterised by a very large reduction in vascular bundle size. Characteristic xylem and phloem cell types were present, but only very small xylem vessels were observed, relative to those found in wild type, *er*f and *px*f lines. Furthermore, tissue layer organisation defects were apparent beyond those previously observed. In particular the clearly defined organisation of endodermal and adjacent phloem cap cells was lacking with the phloem cap appearing to extend into the cortex (Figure 5D) or be absent altogether (Figure 5E). Thus organisation defects occurred out with the vascular cylinder.

**Figure 5.**
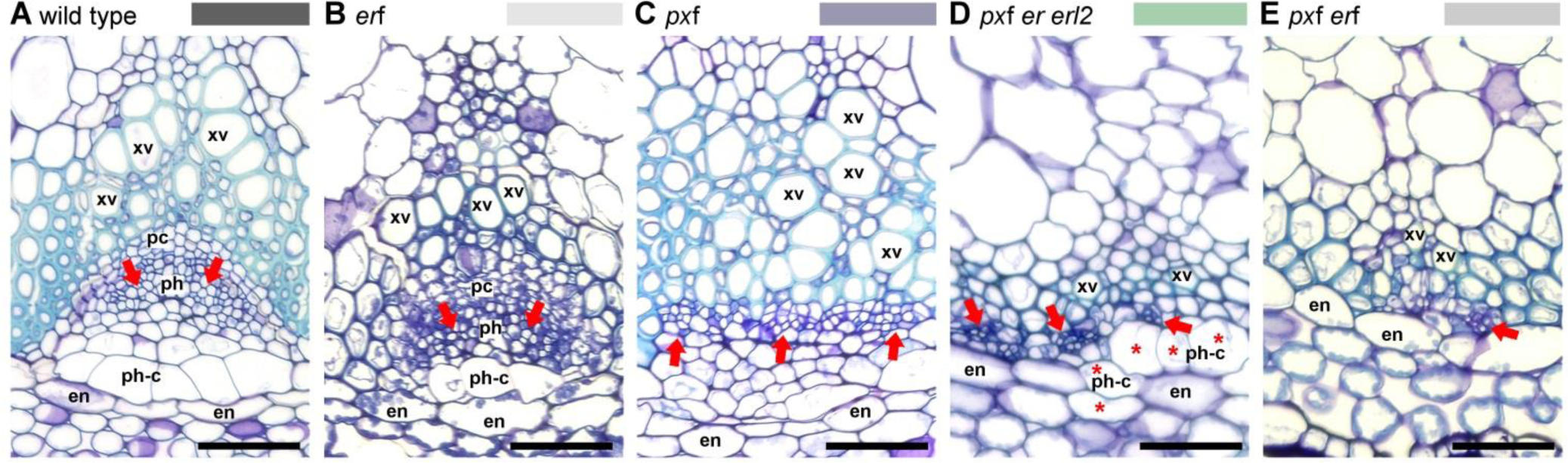
Stem tissue from *px*f *er*f lines. (A) wild type, (B) *er*f, (C) *px*f, (D) *px*f *er erl2*, (E) *px*f *er*f vascular bundles. Phloem arrangement is marked with red arrows. Cells with phloem cap-like morphology are marked with asterisks. Scales are 50 µM; xv is xylem vessel, pc is procambium, ph is phloem, ph-c is phloem cap, en is endodermis.

In hypocotyls, the *px*f *er*f sextuple lines were considerably smaller than all other lines tested (Figure 6). During vascular cylinder development in the embryo, the hypocotyl forms in a diarch pattern with a row of xylem cells that are flanked by two phloem poles (Dolan et al., 1993). As secondary growth proceeds, this organisation develops radial symmetry with phloem present around the circumference of the vascular cylinder (Chaffey et al., 2002). Strikingly, development was perturbed to such a degree in *px*f *er*f mutants that the position of the original phloem poles remained apparent, even after 5 weeks of growth (arrows in figure 6E; Figure S8). This suggests that vascular development was retarded to such a degree that these plants could not make the transformation to true radial growth. In wild type and *er*f lines cell divisions were organised (Figure 6A-B; arrowheads), an aspect of normal vascular development known to perturbed in lines that lack *pxy* and its paralogues (Figure 6C) (Fisher and Turner, 2007). Recent cell divisions were clearly identifiable in the absence of the *PX*f, *ER* and *ERL2* and they remained present, albeit lacking orientation and at a much reduced frequency in *px*f *er*f lines (Figure 6D-E). Thus while not an absolute necessity for formation of either phloem, or xylem vessels, these receptor-kinase families are absolutely essential in specifying their positioning, and coordinating cell division in a manner that allows organised radial expansion and pattern maintenance (Figure 6).

**Figure 6.**
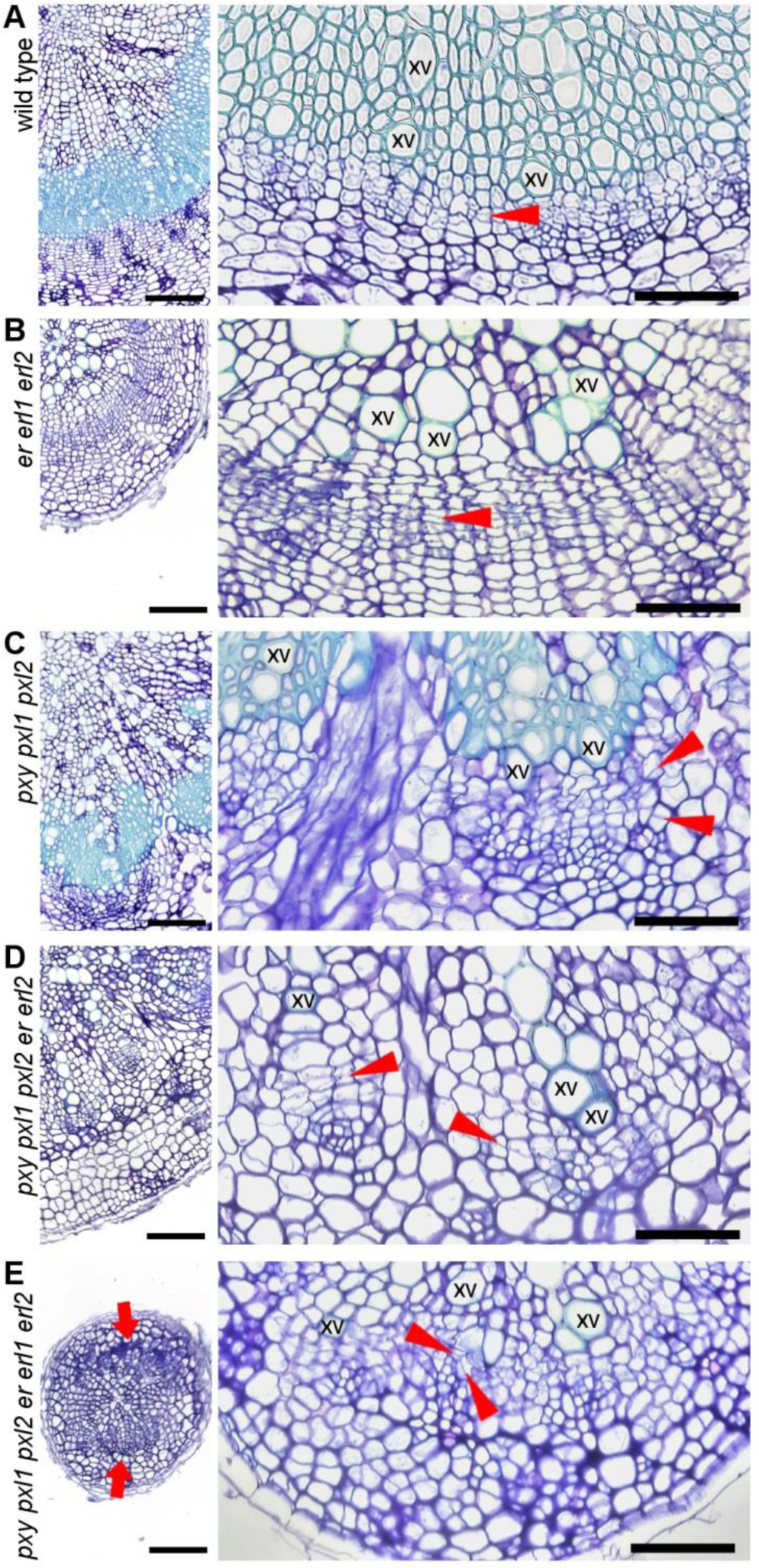
Transverse sections of hypocotyls from *px*f *er*f lines. (A) wild type, (B) *er*f, (C) *px*f, (D) *px*f *er erl2*, (E) *px*f *er*f vascular tissue. Sites of phloem poles in *px*f *er*f are marked with red arrows in left panel of (E). Red arrowheads in panels on right (A-E) align with cell divisions. Scales are 100 µM on left, 50 µM on right; xv is xylem vessel.

### Radial expansion requires crosstalk between tissue layers

An interesting aspect of the interaction between *ER*f and *PX*f signalling is that *ER* ligands are expressed from the endodermis, i.e. from outside the vascular tissues. The observation that *EPFL4* and *EPFL6* expression is perturbed in *px*f *er* mutant stems demonstrates that such coordination occurs. In *Arabidopsis* stems there is very limited radial expansion, but not so in stems of *Nicotiana benthamiana*. Here, interfascicular cambium forms close to the top of the stem, which then expands radially, rather like an *Arabidopsis* hypocotyl, except that endodermal, cortex and epidermal layers are maintained. Thus with this additional level of complexity, fine-tuning of coordinated tissue layer expansion may be more susceptible to perturbation. We generated *Nicotiana benthamiana* lines carrying *AtCLE41*, which encodes for TDIF, under the control of the *35S* promoter (Figure 7). The phenotypes of these constitutive over-expression lines bore similarities to those described in *Arabidopsis* plants and aspen trees carrying similar constructs (Etchells et al., 2015; Etchells and Turner, 2010; Kucukoglu et al., 2017; Strabala et al., 2006). They were characterised by short stature, characteristic organisation defects in vascular tissue, and stem bases with a larger diameter than controls (Figure 7A-D). *35S::AtCLE41 Nicotiana* lines were on average 14.4 mm in diameter compared to 7.6 mm in wild type after 8 weeks of growth. However, in addition to these expected phenotypes, we observed that outer layers of stem tissue had split resulting in prominent lesions in outer tissue layers (Figures 7E-H, see red arrows).

**Figure 7.**
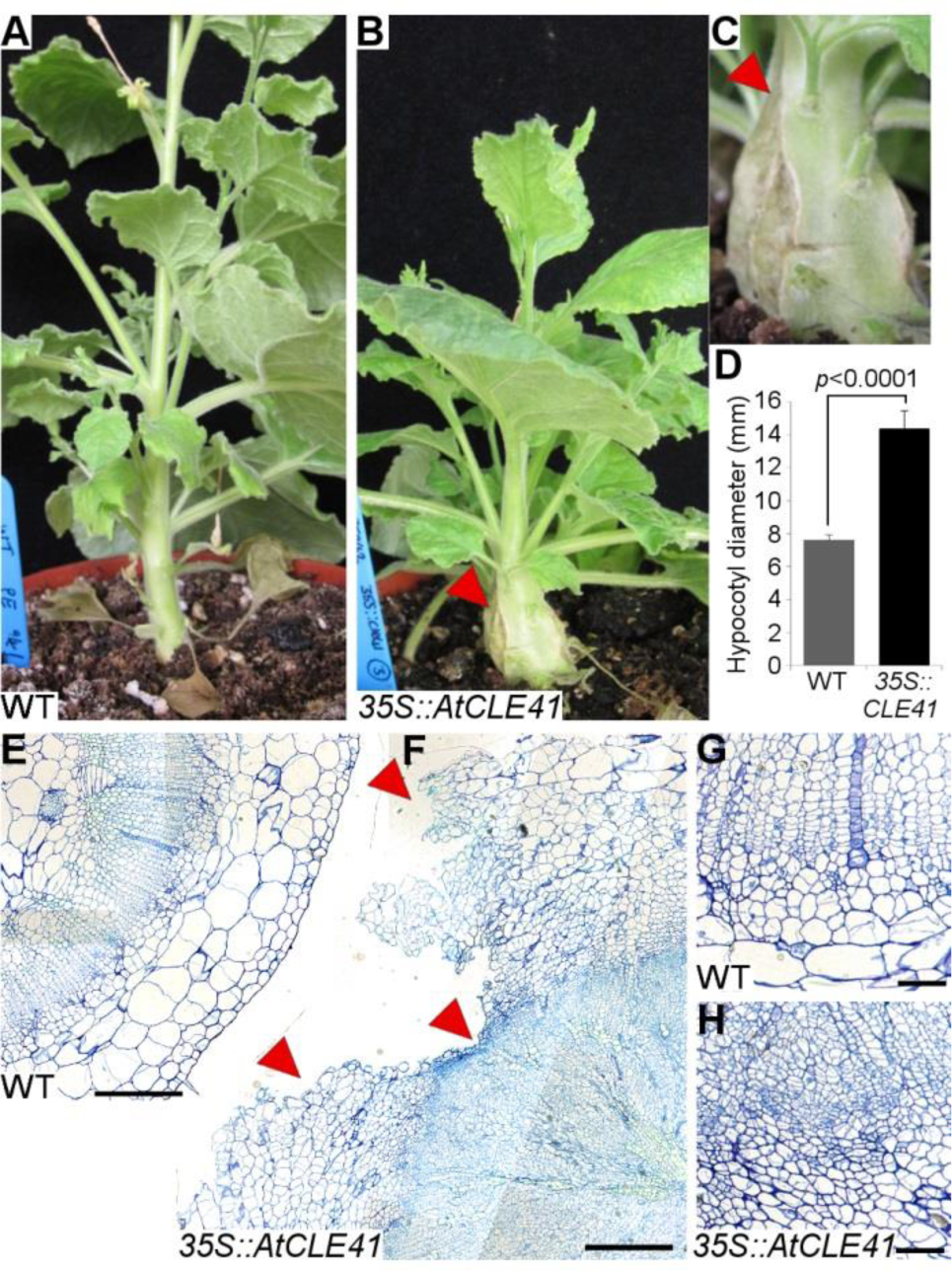
Loss of coordinated expansion in *Nicotiana* lines with disrupted TDIF-PXY signalling. Wild type *Nicotiana benthamiana* plants (A) compared to and *35S::AtCLE41* (B, C) lines showing stem lesions (red arrowhead). (D) Stem diameter of *35S::AtCLE41 Nicotiana* compared to wild type. (E-H) Transverse sections through *Nicotiana* stems. (E) and (F) show areas where lesions are apparent in *35S::AtCLE41*, (I) and (J) show cellular organisation in wild type compared to transgenic lines. Scales are 500 µM (G, H) or 50 µM (I, J).

We tested whether *Nicotiana 35S::AtCLE41* stem lesions were due to large increases in the deposition of vascular tissue by sectioning stems at the site of the split. We observed that the split extended through the outer cell layers, i.e. the epidermis and cortex, but did not extend into the vascular cylinder, thus, in *35S::AtCLE41 Nicotiana* lines the vascular cylinder expands at a greater rate than surrounding tissue, such that lesions develop. This demonstrates that patterning and expansion must be coordinated between the vascular cylinder and outer cell layers for maintenance of tissue integrity during radial growth. It suggests that attenuation of TDIF-PXY signalling may be part of this coordination process. One possibility is that this occurs by ER acting on PXY paralogues.

## Discussion

Plant growth and development requires coordination between expanding tissue layers, particularly where tissue types are organised in concentric rings. Clearly, expansion of inner layers must be coordinated with expansion of outer layers, and our observation that in *Nicotiana*, uncontrolled vascular expansion leads to tissue lesions is a dramatic demonstration that this is the case (Figure 7). Evidence that mechanisms exist to adjust cellular parameters to maintain organisation include the observation that levels of cell expansion differs according to the levels of cell division, such that overall organ size in cell-division mutants is often comparable or, only subtly different to those of wild type plants (De Veylder et al., 2002; Hemerly et al., 1999; Shpak et al., 2004; Ullah et al., 2001). The idea that divisions in one cell layer can influence cell size and organisation in adjacent tissues also comes from experiments where the cell cycle has been manipulated in a cell-type specific manner. Expression of *KRP1* reduces cell division, and when specifically expressed in the epidermal cell layer results in concomitant changes to palisade cell size and density in leaves (Lehmeier et al., 2017).

So how does coordination between tissue layers occur? It was proposed some time ago that the *ER*f could perform this function (Shpak et al., 2004), and this initial suggestion has subsequently been supported by observations that endodermis derived EPFL ligands signal to ER in the phloem to regulate cell division in the adjacent procambium (Uchida et al., 2012; Uchida and Tasaka, 2013)(Figure 1B). Our observation that ER represses *PXL* expression in the stem (Figure 1C, 8A) suggests that endodermis derived signals acting through ER can ultimately attenuate *PX*f-regulated vascular expansion. We also found that the PXf family of receptors, redundantly with *ER*, co-activate expression of *ERL* receptors and their *EPFL* ligands in the stem (Figures 2, S4, 8A). Thus coordination of vascular tissue expansion in stems occurs across multiple tissue layers via a series of feedback loops (Figure 8A). In *Nicotiana 35S::AtCLE41* lines divisions may go unchecked as we found no evidence of regulation of *PXY* by *ER*, so in the presence of very high quantities of TDIF considerable signalling could still occur through PXY.

**Figure 8.**
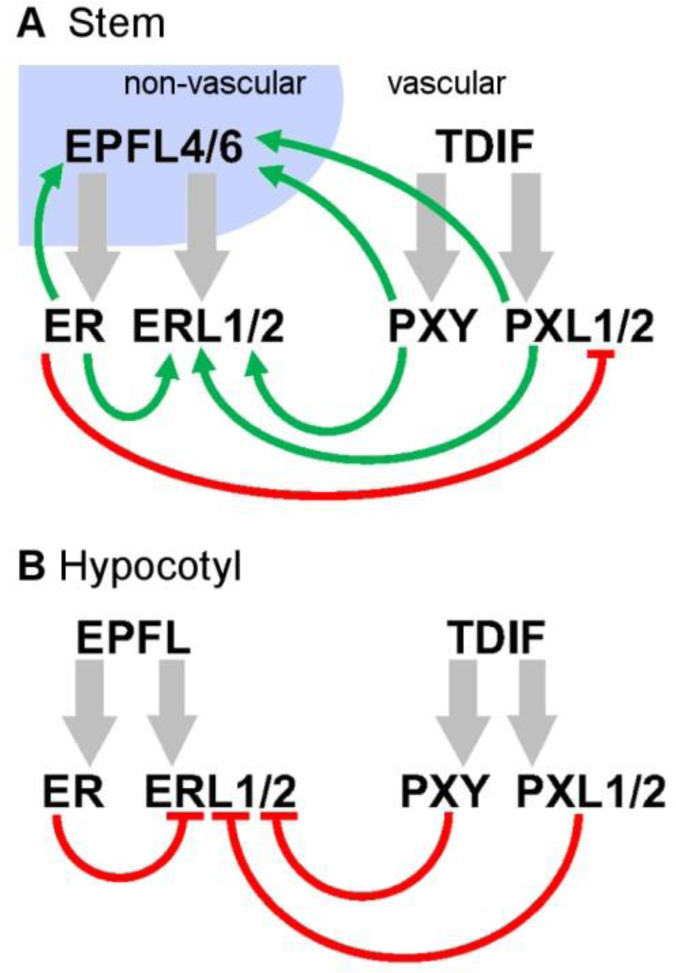
Model showing differences in gene expression regulation in stems and hypocotyls. (A) In the stem, *ER* represses *PXL* gene expression. *PX*f and *ER* act as activators of *ERL* and *EPFL* gene expression. (B) In hypocotyls, negative regulation of *PX*f and *ER* targets predominate. Green arrows show positive influence on gene expression; red blunt ended arrows show repression. Grey arrowsshow ligand-receptor interactions.

In stems, the signalling components that are the focus of this study are expressed in discrete domains, but in the hypocotyl expression patterns of ER and PXY overlap to some extent on the xylem side of the cambium. A direct interaction between these receptors is therefore possible, but a recent global analysis of receptor kinase interactions found no evidence for direct interactions between ERf and PXf family members (Smakowska-Luzan et al., 2018). Our observation that *ERL* expression is de-repressed in the absence of *PX*f and *ER* in hypocotyls (Figure 2) supports the idea that these components interact, at least in part, at the level of gene expression. Perhaps the most striking of our findings was the observation that *ER* and *PX*f regulation of *ERL* expression in the hypocotyl occurred in a manner opposite to that observed in the stem. Here, *ER* and *PX*f combine to repress *ERL* expression, thus while *PX*f and *ER*f are required non-cell autonomously for tissue organisation and expansion in both stems and hypocotyls, the regulatory networks through which development is controlled differ in how they are wired (Figure 8). ERf activity in the epidermis has previously been reported to be buffered by a second receptor, TOO MANY MOUTHS (TMM). Loss of this buffering in *tmm* mutants leads to opposite stomatal spacing phenotypes in spatially separate cotyledon, where stomata cluster, compared to hypocotyls where stomata are absent. Differing ligand availability in cotyledon and hypocotyl is thought to account for this difference (Abrash et al., 2011). While EPFL4 and 6 have been demonstrated to act as ERf ligands in the inner tissues in stems, less is known about ligand expression pattern in hypocotyls. It remains to be determined whether the difference in ERL regulation by *ER* and *PX*f in stem and hypocotyl could be due to differing complements of co-receptors and ligands in these differing locations.

In both stems and hypocotyls, tissue layers are arranged largely in concentric rings. However, in *Arabidopsis*, stem and hypocotyl differ in that the hypocotyl undergoes radial growth, but the vast majority of the stem does not. Radial hypocotyl growth is largely the consequence of expansion of a pattern that is laid down in the embryo, but in stems, *de novo* patterning must occur below the shoot apical meristem. In stems the epidermis, cortex and endodermal layers are maintained, but in hypocotyls they are lost. Nevertheless in both stem and hypocotyl, the xylem, (pro)cambium and phloem must be specified in adjacent tissue layers in a coordinated manner. Our mutant analysis demonstrates that *PX*f and *ER*f are central to maintaining this organisation (Figures 1, 5–6). The result of loss of this signalling, as determined by analysis of *px*f *er* quadruple mutants is severe disruption to vascular pattern such that in stems, vascular tissue is no longer found in discrete bundles, but scattered around the stem adjacent to the endodermis (Figure 2). Removal of *PX*f and *ER*f families in hypocotyls results in prominent proliferation defects (Figure 6), but perhaps significantly, the ability to adjust cell size to compensate for the profound reductions in cell division (Figure 4) was also lost. This is in contrast to the consequences of losing the *ERECTA* family alone, as cell size adjustments are a feature of *er*f mutants (Shpak et al., 2004). Thus these observations support the idea that one function of the interaction between *ER*f and *PX*f is coordination of tissue expansion. We propose that with these signalling mechanisms removed, the positional information that must be interpreted for cell morphology adjustments to occur is missing.

In hypocotyls, the switch from primary to secondary growth is relatively unstudied, as are the events that occur in the rib zone below the shoot apical meristem where stem vascular tissues are formed. However, oriented cell divisions and the development of organ boundaries in the rib zone have been reported to be regulated by a homeodomain transcription factor, REPLUMLESS (RPL). Pertinent to the results obtained here, RPL was found to occupy the promoters of *PXY*, *CLE41*, *CLE42*, *ER*, *ERL1*, *ERL2*, and *CHAL* in ChIP-Seq experiments (Bencivenga et al., 2016). RPL is localised to the cytoplasm unless present in a heterodimer with class I KNOX protein, such as BREVIPEDICELLUS (Bhatt et al., 2004). *rpl bp* double mutants, particularly those in the L*er* background that lacks a functional copy of *ER*, demonstrate considerable defects in vascular development (Etchells et al., 2012; Smith and Hake, 2003). Thus events in the rib zone, controlled by RPL could set up the initial pattern. Our genetic analysis demonstrates that however the pattern is initiated, it is maintained by interacting signalling pathways characterised by members of the ERECTA and PXY families. Such overlapping signals may be involved in coordinating growth in adjacent tissues in other developmental contexts.

## Acknowledgements

We thank Miguel de Lucas, Keith Lindsey, and Jen Topping for critical reading of the manuscript. The authors are grateful to Keiko Torii for sharing *er* and *erl* mutants, and *ER*f reporter lines, and to the Nottingham Arabidopsis Stock Centre for providing other genetic resources. This work was funded by the European Union (project number 329978 - Marie Skladowska Curie Fellowship to JPE) and the BBSRC (grant number BB/H019928 to JPE and SRT and an NLD-DTP studentship to KSB, JPE and IHJ). The authors gratefully acknowledge a travel grant from Henan Agricultural University to NW.

## Literature Cited

Abrash, E. B., Davies, K. A. and Bergmann, D. C. (2011). Generation of Signaling Specificity in Arabidopsis by Spatially Restricted Buffering of Ligand–Receptor Interactions. Plant Cell 23, 2864–2879.

Bae, S., Park, J. and Kim, J.-S. (2014). Cas-OFFinder: a fast and versatile algorithm that searches for potential off-target sites of Cas9 RNA-guided endonucleases. Bioinformatics 30, 1473–1475.

Bemis, S. M., Lee, J. S., Shpak, E. D. and Torii, K. U. (2013). Regulation of floral patterning and organ identity by Arabidopsis ERECTA-family receptor kinase genes. J Exp Bot 64, 5323–5333.

Bencivenga, S., Serrano-Mislata, A., Bush, M., Fox, S. and Sablowski, R. (2016). Control of Oriented Tissue Growth through Repression of Organ Boundary Genes Promotes Stem Morphogenesis. Developmental Cell 39, 198–208.

Bhatt, A. M., Etchells, J. P., Canales, C., Lagodienko, A. and Dickinson, H. (2004). VAAMANA--a BEL1-like homeodomain protein, interacts with KNOX proteins BP and STM and regulates inflorescence stem growth in Arabidopsis. Gene 328, 103–111.

Chaffey, N., Cholewa, E., Regan, S. and Sundberg, B. (2002). Secondary xylem development in Arabidopsis: a model for wood formation. Physiol Plant 114, 594–600.

Clough, S. J. and Bent, A. F. (1998). Floral dip: a simplified method for Agrobacterium-mediated transformation of Arabidopsis thaliana. Plant Journal 16, 735–743.

De Veylder, L., Beeckman, T., Beemster, G. T. S., Engler, J. D., Ormenese, S., Maes, S., Naudts, M., Van der Schueren, E., Jacqmard, A., Engler, G., et al. (2002). Control of proliferation, endoreduplication and differentiation by the Arabidopsis E2Fa-DPa transcription factor. Embo J 21, 1360–1368.

Dolan, L., Janmaat, K., Willemsen, V., Linstead, P., Poethig, S., Roberts, K. and Scheres, B. (1993). CELLULAR-ORGANIZATION OF THE ARABIDOPSIS-THALIANA ROOT. Development 119, 71–84.

Efroni, I., Mello, A., Nawy, T., Ip, P.-L., Rahni, R., DelRose, N., Powers, A., Satija, R. and Birnbaum, K. D. (2016). Root Regeneration Triggers an Embryo-like Sequence Guided by Hormonal Interactions. Cell 165, 1721–1733.

Etchells, J. P., Mishra, Laxmi S., Kumar, M., Campbell, L. and Turner, Simon R. (2015). Wood Formation in Trees Is Increased by Manipulating PXY-Regulated Cell Division. Current Biology 25, 1050–1055.

Etchells, J. P., Moore, L., Jiang, W. Z., Prescott, H., Capper, R., Saunders, N. J., Bhatt, A. M. and Dickinson, H. G. (2012). A role for BELLRINGER in cell wall development is supported by loss-of-function phenotypes. BMC Plant Biol 12, 212.

Etchells, J. P., Provost, C. M., Mishra, L. and Turner, S. R. (2013). WOX4 and WOX14 act downstream of the PXY receptor kinase to regulate plant vascular proliferation independently of any role in vascular organisation. Development 140, 2224–2234.

Etchells, J. P. and Turner, S. R. (2010). The PXY-CLE41 receptor ligand pair defines a multifunctional pathway that controls the rate and orientation of vascular cell division. Development 137, 767–774.

Feng, X. and Dickinson, H. G. (2010). Tapetal cell fate, lineage and proliferation in the Arabidopsis anther. Development 137, 2409–2416.

Fisher, K. and Turner, S. (2007). PXY, a receptor-like kinase essential for maintaining polarity during plant vascular-tissue development. Current Biology 17, 1061–1066.

Hemerly, A. S., Ferreira, P. C. G., Van Montagu, M. and Inze, D. (1999). Cell cycle control and plant morphogenesis: is there an essential link? Bioessays 21, 29–37.

Hirakawa, Y., Shinohara, H., Kondo, Y., Inoue, A., Nakanomyo, I., Ogawa, M., Sawa, S., Ohashi-Ito, K., Matsubayashi, Y. and Fukuda, H. (2008). Non-cell-autonomous control of vascular stem cell fate by a CLE peptide/receptor system. Proceedings of the National Academy of Sciences, USA 105, 15208–15213.

Horiguchi, G. and Tsukaya, H. (2011). Organ Size Regulation in Plants: Insights from Compensation. Frontiers in Plant Science 2.

Horsch, R. B., Fry, J. E., Hoffmann, N. L., Eichholtz, D., Rogers, S. G. and Fraley, R. T. (1985). A Simple and General Method for Transferring Genes into Plants. Science 227, 1229–1231.

Ikematsu, S., Tasaka, M., Torii, K. U. and Uchida, N. (2017). ERECTA-family receptor kinase genes redundantly prevent premature progression of secondary growth in the Arabidopsis hypocotyl. New Phytologist 213, 1697–1709.

Ito, Y., Nakanomyo, I., Motose, H., Iwamoto, K., Sawa, S., Dohmae, N. and Fukuda, H. (2006). Dodeca-CLE peptides as suppressors of plant stem cell differentiation. Science 313, 842–845.

Jia, G., Liu, X., Owen, H. A. and Zhao, D. (2008). Signaling of cell fate determination by the TPD1 small protein and EMS1 receptor kinase. Proceedings of the National Academy of Sciences 105, 2220–2225.

Kajala, K., Ramakrishna, P., Fisher, A., C. Bergmann, D., De Smet, I., Sozzani, R., Weijers, D. and Brady, S. M. (2014). Omics and modelling approaches for understanding regulation of asymmetric cell divisions in arabidopsis and other angiosperm plants. Annals of Botany 113, 1083–1105.

Kimura, Y., Tasaka, M., Torii, K. U. and Uchida, N. (2018). ERECTA-family genes coordinate stem cell functions between the epidermal and internal layers of the shoot apical meristem. Development 145.

Kucukoglu, M., Nilsson, J., Zheng, B., Chaabouni, S. and Nilsson, O. (2017). WUSCHEL-RELATED HOMEOBOX4 (WOX4)-like genes regulate cambial cell division activity and secondary growth in Populus trees. New Phytologist 215, 642–657.

Lehmeier, C., Pajor, R., Lundgren Marjorie, R., Mathers, A., Sloan, J., Bauch, M., Mitchell, A., Bellasio, C., Green, A., Bouyer, D., et al. (2017). Cell density and airspace patterning in the leaf can be manipulated to increase leaf photosynthetic capacity. The Plant Journal 92, 981–994.

Ragni, L., Nieminen, K., Pacheco-Villalobos, D., Sibout, R., Schwechheimer, C. and Hardtke, C. S. (2011). Mobile Gibberellin Directly Stimulates Arabidopsis Hypocotyl Xylem Expansion. Plant Cell 23, 1322–1336.

Ramakers, C., Ruijter, J. M., Deprez, R. H. L. and Moorman, A. F. M. (2003). Assumption-free analysis of quantitative real-time polymerase chain reaction (PCR) data. Neuroscience Letters 339, 62–66.

Shpak, E. D., Berthiaume, C. T., Hill, E. J. and Torii, K. U. (2004). Synergistic interaction of three ERECTA-family receptor-like kinases controls Arabidopsis organ growth and flower development by promoting cell proliferation. Development 131, 1491–1501.

Shpak, E. D., Lakeman, M. B. and Torii, K. U. (2003). Dominant-negative receptor uncovers redundancy in the Arabidopsis ERECTA leucine-rich repeat receptor-like kinase signaling pathway that regulates organ shape. Plant Cell 15, 1095–1110.

Shpak, E. D., McAbee, J. M., Pillitteri, L. J. and Torii, K. U. (2005). Stomatal Patterning and Differentiation by Synergistic Interactions of Receptor Kinases. Science 309, 290–293.

Smakowska-Luzan, E., Mott, G. A., Parys, K., Stegmann, M., Howton, T. C., Layeghifard, M., Neuhold, J., Lehner, A., Kong, J., Grünwald, K., et al. (2018). An extracellular network of Arabidopsis leucine-rich repeat receptor kinases. Nature 553, 342.

Smith, H. M. S. and Hake, S. (2003). The Interaction of Two Homeobox Genes, BREVIPEDICELLUS and PENNYWISE, Regulates Internode Patterning in the Arabidopsis Inflorescence. Plant Cell 15, 1717–1727.

Strabala, T. J., O’Donnell, P. J., Smit, A. M., Ampomah-Dwamena, C., Martin, E. J., Netzler, N., Nieuwenhuizen, N. J., Quinn, B. D., Foote, H. C. C. and Hudson, K. R. (2006). Gain-of-function phenotypes of many CLAVATA3/ESR genes, including four new family members, correlate with tandem variations in the conserved CLAVATA3/ESR domain. Plant Physiology 140, 1331–1344.

ten Hove, C. A., Lu, K.-J. and Weijers, D. (2015). Building a plant: cell fate specification in the early Arabidopsis embryo. Development 142, 420–430.

Torii, K. U., Mitsukawa, N., Oosumi, T., Matsuura, Y., Yokoyama, R., Whittier, R. F. and Komeda, Y. (1996). The arabidopsis ERECTA gene encodes a putative receptor protein kinase with extracellular leucine-rich repeats. Plant Cell 8, 735–746.

Uchida, N., Lee, J. S., Horst, R. J., Lai, H.-H., Kajita, R., Kakimoto, T., Tasaka, M. and Torii, K. U. (2012). Regulation of inflorescence architecture by intertissue layer ligand–receptor communication between endodermis and phloem. Proc Natl Acad Sci USA.

Uchida, N., Shimada, M. and Tasaka, M. (2013). ERECTA-Family Receptor Kinases Regulate Stem Cell Homeostasis via Buffering its Cytokinin Responsiveness in the Shoot Apical Meristem. Plant Cell Physiol 54, 343–351.

Uchida, N. and Tasaka, M. (2013). Regulation of plant vascular stem cells by endodermis-derived EPFL-family peptide hormones and phloem-expressed ERECTA-family receptor kinases. J Exp Bot.

Ullah, H., Chen, J.-G., Young, J. C., Im, K.-H., Sussman, M. R. and Jones, A. M. (2001). Modulation of Cell Proliferation by Heterotrimeric G Protein in <em>Arabidopsis</em>. Science 292, 2066–2069.

Wang, Z.-P., Xing, H.-L., Dong, L., Zhang, H.-Y., Han, C.-Y., Wang, X.-C. and Chen, Q.-J. (2015). Egg cell-specific promoter-controlled CRISPR/Cas9 efficiently generates homozygous mutants for multiple target genes in Arabidopsis in a single generation. Genome Biology 16, 144.

Wunderling, A., Ripper, D., Barra-Jimenez, A., Mahn, S., Sajak, K., Targem, M. B. and Ragni, L. (2018). A molecular framework to study periderm formation in Arabidopsis. New Phytologist 0.

Xie, K., Zhang, J. and Yang, Y. (2014). Genome-Wide Prediction of Highly Specific Guide RNA Spacers for CRISPR–Cas9-Mediated Genome Editing in Model Plants and Major Crops. Mol Plant 7, 923–926.

Xing, H.-L., Dong, L., Wang, Z.-P., Zhang, H.-Y., Han, C.-Y., Liu, B., Wang, X.-C. and Chen, Q.-J. (2014). A CRISPR/Cas9 toolkit for multiplex genome editing in plants. BMC Plant Biol 14, 327.

